# Blue Light Negatively Regulates Tolerance to Phosphate Deficiency in Arabidopsis

**DOI:** 10.1101/235952

**Authors:** Chuan-Ming Yeh, Koichi Kobayashi, Sho Fujii, Hidehiro Fukaki, Nobutaka Mitsuda, Masaru Ohme-Takagi

## Abstract

Plants have evolved mechanisms to improve utilization efficiency or acquisition of inorganic phosphate (Pi) in response to Pi deficiency, such as altering root architecture, secreting acid phosphatases, and activating the expression of genes related to Pi uptake and recycling. Although many genes responsive to Pi starvation have been identified, transcription factors that affect tolerance to Pi deficiency have not been well characterized. We show here that defect in the *ELONGATED HYPOCOTYL 5* (*HY5*) transcription factor gene results in tolerance to Pi deficiency in Arabidopsis. The primary root length of *hy5* was only slightly inhibited under Pi deficient condition and its fresh weight was significantly higher than that of wild type. The Pi deficiency-tolerant phenotype of *hy5* was similarly observed when grown on the medium without Pi. In addition, a double mutant, *hy5slr1,* without lateral roots also showed tolerance to phosphate deficiency, indicating that the tolerance of *hy5* does not result from increase of external Pi uptake and may be related to internal Pi utilization or recycling. Moreover, we found that blue light negatively regulates tolerance to Pi-deficiency and that *hy5* exhibits tolerance to Pi deficiency due to blockage of blue-light responses. Collectively, this study points out light quality may play an important role in the regulation of internal Pi recycling and utilization efficiency. Also, it may contribute to reducing Pi fertilizer requirements in plants through a proper illumination.

## Introduction

Inorganic phosphate (Pi) is an essential constituent of ATP, nucleic acids and membrane phospholipids. In addition, it is crucial to various cellular metabolic pathways, including photosynthesis, glycolysis, respiration, signal transduction and carbohydrate metabolism (Ticconi AND Abel 2004, Péret et al. 2011, Niu 2013). However, Pi is easily chelated by soil particles or formed insoluble complexes with aluminum or iron at acid pH and with calcium at alkaline pH leading to a low mobility and availability in soils (Wissuwa 2003, Gaxiola et al. 2011). Therefore, available soil Pi concentrations are often less than the requirement for optimal crop production (Nussaume et al. 2011, Péret et al. 2011, Niu 2013). Plants have evolved adaptive mechanisms to acquire and recycle Pi in response to Pi deficiency. Alteration of root architecture, such as enhancement of lateral root growth and root hair formation, increases root surface areas for Pi absorption (Ticconi AND Abel 2004, Péret et al. 2011). Induction of high-affinity Pi transporter genes increases uptake of soluble Pi, while activation or secretion of acid phosphatases, ribonucleases, and organic acids enhances scavenging of extracellular Pi from insoluble organic complexes. In addition, the activities of acid phosphatases and ribonucleases also help release Pi from intracellular organic Pi-containing molecules (Raghothama 2000, Poirier and Bucher 2002, Nussaume et al. 2011). To improve Pi use efficiency, plants substitute bypass pathways that do not require Pi for metabolic processes requiring Pi (Plaxton and Tran 2011). Replacing membrane phospholipids with non-P-containing glycolipids also plays an important role in the supply of free Pi during Pi deficiency (Kobayashi et al. 2006).

Many efforts have been made to unravel the molecular mechanisms that regulate Pi starvation responses (PSRs). An array of Pi starvation-induced (PSI) genes have been identified by transcriptome studies (Wu et al. 2003, Misson et al. 2005, Thibaud et al. 2010, Woo et al. 2012) and a series of *hypersensitive to phosphate starvation* (*hps*) mutants have been isolated and characterized (Yeh et al. 2017). Although various plant transcription factors (TFs) affect PSRs, the transcriptional regulation of these processes is not yet well elucidated. *AtPHR1* (*PHOSPHATE STARVATION RESPONSE 1*) is the first Arabidopsis TF gene shown to mediate diverse PSRs (Rubio et al. 2001). Although *AtPHR1* is not Pi starvation-inducible, PHR1 regulates a subset of PSI genes through the miR399-PHO2 (an ubiquitin-conjugating E2 enzyme) signaling pathway (Bari et al. 2006, Chiou et al. 2006). *AtPHR1*, *AtPHL1* (*PHR1-like 1*), and their two rice orthologues, *OsPHR1* and *OsPHR2,* have been identified as having partially redundant functions (Zhou et al. 2008, Bustos et al. 2010, Liu et al. 2010). In addition, several TFs have been identified as negative regulators of PSRs in Arabidopsis. BHLH32, a basic helix-loop-helix TF, negatively regulates anthocyanin accumulation, root hair formation, and induction of the PSI genes (Chen et al. 2007). *AtMYB62* is low-Pi-inducible and mediates its negative effects on PSRs through modulation of gibberellin metabolism (Devaiah et al. 2009). WRKY6 and WRKY42 negatively regulate the expression of *PHOSPHATE1* (*PHO1*), which is responsible for Pi translocation from root to shoot in Arabidopsis (Hamburger et al. 2002, Chen et al. 2009). AtWRKY75 and AtZAT6 have been reported to regulate root development and Pi acquisition, although they may not be specific to PSRs due to their responsiveness to multiple nutrient deficiencies (Devaiah et al. 2007a and 2007b). In recent years, several Arabidopsis TF genes, such as *AtERF070, APSR1, AtMYB2* and *AL6*, have been shown to be involved in the regulation of root growth and architecture under Pi deficiency (Yeh and Ohme-Takagi 2015).

Adding Pi fertilizer can improve soil Pi levels; however, the world’s Pi rock reserves may be exhausted within 120 years (Gilbert 2009; Nussaume et al. 2011) and the demand for Pi fertilizers will likely increase to support crop productivity for the growing global population (Nussaume et al. 2011, Péret et al. 2011). In addition, the low solubility of Pi in soils often causes over-application of chemical fertilizers, subsequently, leading to potential threats to the environment and the ecosystem (Gaxiola et al. 2011, Péret et al. 2011). Therefore, proper utilization of the remaining Pi reserves is important to reduce Pi resource depletion and environmental threaten. To this end, development of crops with tolerance to Pi deficiency is required, especially if crops can be manipulated to possess higher ability for Pi recycling or Pi utilization efficiency.

In this study, we identified a Pi deficiency-tolerant *hy5-215* mutant with defect in the Arabidopsis bZIP TF ELONGATED HYPOCOTYL 5 (HY5). Under Pi-deficient conditions, primary root length and seedling fresh weight were reduced to a lesser extent in the *hy5-215* mutant compared to the wild type (WT). The Pi-deficiency tolerance phenotype of *hy5-215* did not change in plants grown on medium without Pi, indicating that this tolerance may be related to an enhanced internal Pi utilization but not uptake of external Pi. Furthermore, we found that continuous blue light accelerate sensitivity to Pi deficiency in WT and elimination from blue light improve WT tolerance to Pi deficiency. Our results indicate that blue light plays a negative role in Pi deficiency tolerance and *hy5-215* exhibits tolerance to Pi deficiency probably due to blockage of blue-light responses.

## Results and Discussion

### Tolerant phenotypes of *hy5-215* mutants under Pi deficiency

To identify transcription factors (TFs) that can be manipulated to allow plants growing well under minimal Pi fertilization, we grew Arabidopsis mutants in Pi-deficient conditions and screened for plant phenotypes indicative of tolerance to Pi deficiency: larger plant size, longer primary root (PR) length, and lower anthocyanin accumulation than wild type (WT). The *hy5-215* mutant with a defect in *HY5,* which encodes a bZIP TF that functions in photopmophogenesis, exhibited a Pi deficiency-tolerant phenotype. The PR lengths of WT were significantly reduced under Pi-deficient conditions (10 μM Pi) when compared with those grown under Pi-sufficient conditions (625 μM Pi) while only slight inhibition of PR growth was observed in the *hy5-215* mutant between Pi-sufficient and Pi-deficient conditions (Fig. 1A, B). WT fresh weight declined to 37% under Pi deficient-conditions compared to Pi-sufficient conditions while *hy5-215* fresh weight declined to 65% under Pi deficient-conditions compared to Pi-sufficient conditions (Fig. 1C and Supplementary Fig. S1). We also confirmed the tolerance of *hy5-215* to Pi deficiency by examination of several well-known PSRs including expression of ribonuclease, purple acid phosphatase and anthocyanin biosynthesis genes (Supplementary Note 1 and Supplementary Fig. S2-4).

**Figure 1.**
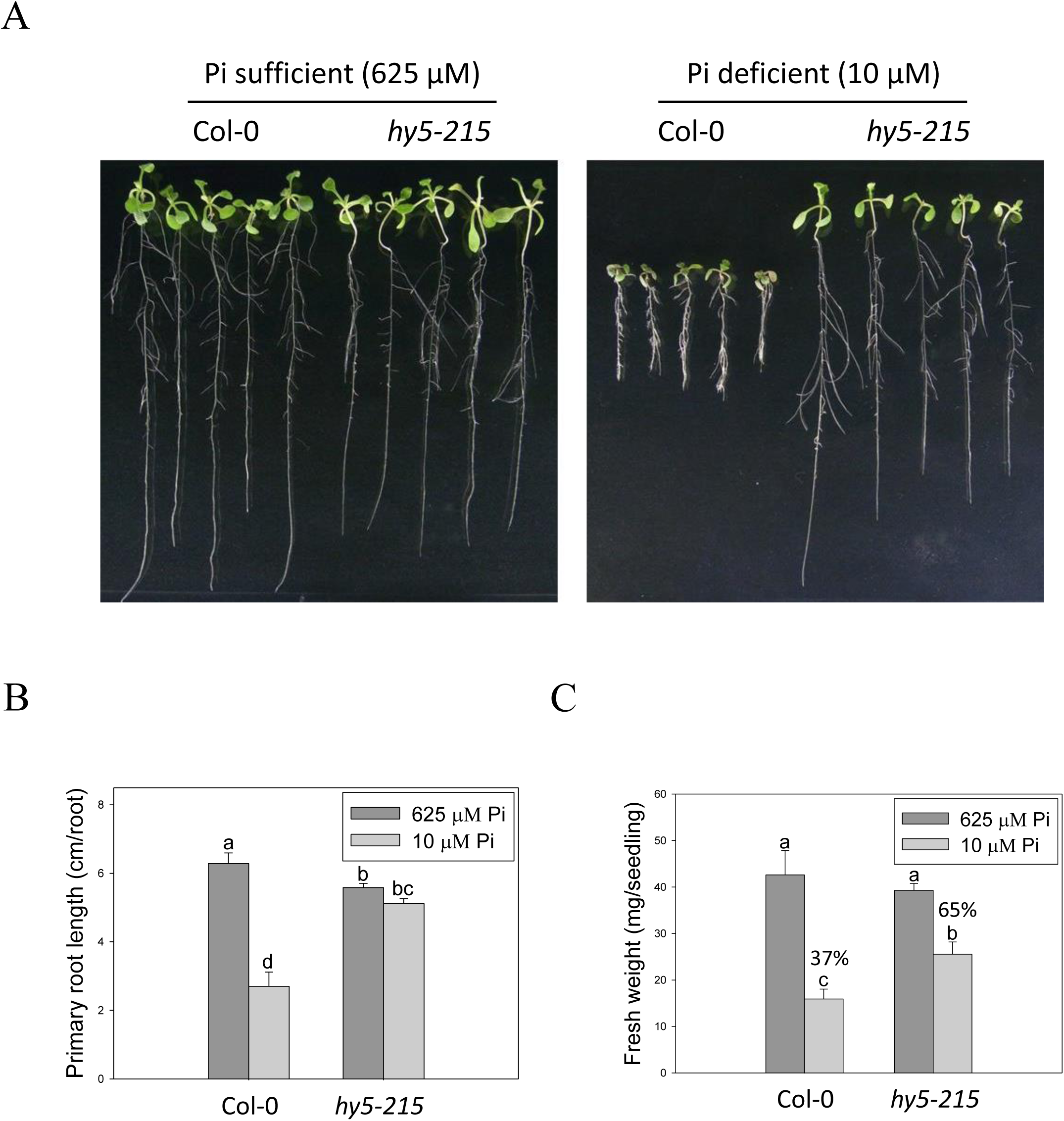
Primary root length and fresh weight of wild-type and mutant seedlings in response to Pi treatment. (A) Wildtype (Col-0) and *hy5-215* seedlings grown in Pi-sufficient (625 μM Pi) and Pi-deficient (10 μM Pi) conditions. (B) Primary root (PR) lengths after growth on vertical plates for 10 days. (C) Seedling fresh weights after growth on horizontal plates for 14 days. Data represent the means ± standard error (SE) of four independent experiments. Different letters above the bars indicate statistically significant differences among the means based on ANOVA (Analysis of Variance) followed by Fisher’s LSD (Least Significant Difference) tests (*P* <0.05).

### Alteration of root architecture in *hy5-215* is not responsible to Pi-deficiency tolerance

Plant root architecture, the spatial arrangement of a root system, is highly plastic in response to depletion of mineral nutrients. Modifications of RA through altering the number, length, angle and diameter of roots or root hairs enable plants to cope with nutrient shortages (Gruber et al. 2013). The “topsoil foraging” strategy is employed to get immobile Pi from the Pi-enriched upper-layer soil under Pi deficiency; in topsoil foraging, plants inhibit PR growth but enhance lateral root (LR) growth and root hair formation, thus increasing the surface area available for Pi uptake (Péret et al. 2011, Sato and Miura 2011, Niu 2013). In this study, a great number of root hairs were initiated in the WT under Pi-deficient conditions, whereas *hy5-215* formed fewer and shorter root hairs (Fig. 2A), suggesting that *hy5-215* may not show as strong of a response to Pi deficiency as WT. However, LR numbers and lengths were not enhanced by low-Pi treatment in both WT and *hy5-215*. Instead, LR growth was repressed by our Pi deficiency condition (Fig. 2B-D). This inconsistency may result from different Pi concentrations and experimental conditions used in the different studies. Plants grown at relatively higher levels of Pi (> 1 mM) in Pi-sufficient media form fewer or almost no LRs (Pérez-Torres et al. 2008, Lei et al. 2011). However, Pi-sufficient treatment (625 μM) in this work induces much more LR formation and growth. This is in agreement with some previous reports that use relative lower concentrations for Pi-sufficient treatments (Devaiah et al. 2007a, Pérez-Torres et al. 2008, Devaiah et al. 2009, Lei et al. 2011, Gruber et al. 2013).

**Figure 2.**
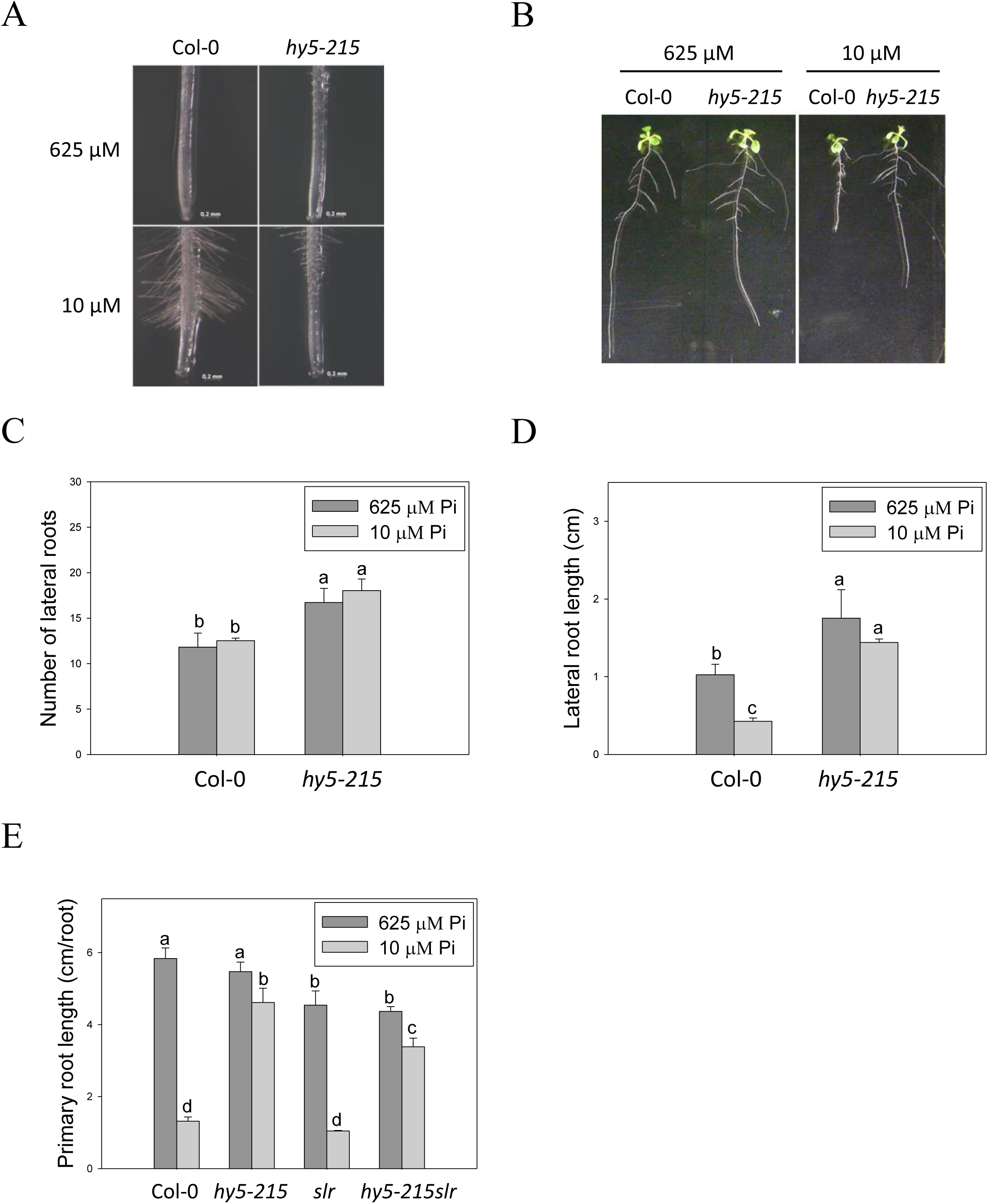
Root hair formation and root architecture of wild-type and mutant seedlings in response to Pi treatment. (A) Root hair formation of Col-0 and *hy5-215* after growth of 7 days. (B) Root architecture of Col-0 and *hy5-215* after growth of 10 days. (C) Increase of LR number in *hy5-215* plants. (D) Increase of LR length in *hy5-215* plants. (E) PR length in Col-0, *hy5-215, slr,* and *hy5-215slr-1.* All the seedlings were grown on 1/2 MS medium with 625 or 10 μM Pi for 7 to 10 days. Data represent means ± SE of four independent experiments. Different letters above the bars indicate statistically significant differences among the means based on ANOVA followed by Fisher’s LSD tests (*P* <0.05).

Although LR growth was not enhanced by Pi starvation in this study, a root system possessing more and longer LRs was found in *hy5-215* in both Pi-sufficient and Pi-deficient conditions (Fig. 2B-D). To examine whether the increased LR number and lengths contribute to the Pi-deficiency tolerance in *hy5-215*, a double mutant constructed with *hy5-215* and *solitary-root-1* (*slr-1*), a gain-of-function mutant of IAA14 (a repressor of auxin signaling) that produces no LRs, was examined under Pi deficiency (Fukaki et al. 2002; Kobayashi et al. 2012). The *hy5-215 slr-1* double mutant showed a long-hypocotyl phenotype similar to that of *hy5-215* and a PR lacking LR growth similar to the *slr-1* phenotype (Fig. 2E). Interestingly, the PR elongation of *hy5-215 slr-1* seedlings was only slightly inhibited by Pi deficiency, although the PR of *hy5-215 slr-1* was shorter than that of *hy5-215* in the respective conditions. The results revealed that LR growth is beneficial for growth on Pi-deficient medium, but the change in *hy5-215* root architecture does not appear to be responsible for the observed tolerance to Pi deficiency in *hy5-215.* Auxin signaling was reported to be enhanced in Arabidopsis *hy5* mutants (Oyama et al. 1997, Cluis et al. 2004), whereas it may be repressed in *hy5-215 slr-1* mutants due to the gain-of-function mutation of *SLR/IAA14.* Therefore, the similar tolerance phenotypes between *hy5-215 slr-1* and *hy5-215* also suggest that auxin signaling may not be responsible for the Pi-deficiency tolerance in *hy5-215.*

### External Pi acquisition is not involved in Pi-deficiency tolerance of *hy5-215*

Enhancement of Pi influx through induction of high-affinity Pi transporter genes is one of the conserved strategies evolved by plants to optimize their growth in response to Pi limitation. There are nine genes encoding *PHT* homologs (*PHT1;1–PHT1;9*) in the Arabidopsis genome. Most of the *PHT1* family genes are strongly induced by low Pi treatment within the first 12 hours (Bayle et al. 2011, Nussaume et al. 2011). Functional studies show a major role for PHT1 in Pi acquisition in roots from Pi-deficient environment; however, some of the PHTs are also required for Pi mobilization (PHT1;5), flower development (PHT1;6) and Pi uptake in Pi replete condition (PHT1;1 and PHT1;4) (Nussaume et al. 2011, Nagarajan et al. 2011). In this study, we found that expression of *PHT1* genes was lower in *hy5-215* shoots than in the WT, suggesting *hy5-215* may not be as deficient as WT under low Pi treatment (Supplementary Fig. S5). However, several *PHT1* genes were induced in a higher level in *hy5-215* roots under both sufficient and deficient conditions (Supplementary Table S1). To demonstrate whether the higher *PHT1* gene expression in *hy5-215* roots can increase Pi uptake and subsequently contributes to Pi-deficiency tolerance, the free Pi content were measured. A great reduction of Pi level was found in *hy5-215* shoots under Pi sufficient condition, although Pi content was slightly higher in *hy5-215* shoots than in WT shoots under Pi deficiency (Fig. 3A). There was no significant difference between WT and *hy5-215* in roots (Fig. 3B). The results indicated that the elevated amounts of *PHT1* transcripts in *hy5-215* roots might not or only partially contribute to Pi deficiency tolerance of *hy5-215.* To verify this finding, we cultured WT and *hy5-215* plants on Pi-free media. The *hy5-215* plants exhibited similar growth on Pi-free medium and on Pi-deficient medium containing 10 μM Pi. The PR length of *hy5-215* grown on Pi-free medium was only slightly diminished compared to that of plants grown on Pi-deficient medium (Fig. 3C). Altogether, these results indicated that the tolerance of *hy5-215* to Pi deficiency was not related to extracellular Pi acquisition. Furthermore, it also suggested the pre-accumulated Pi in seeds during seed development is sufficient to support *hy5-215* growth at the early stages of Pi deficiency.

**Figure 3.**
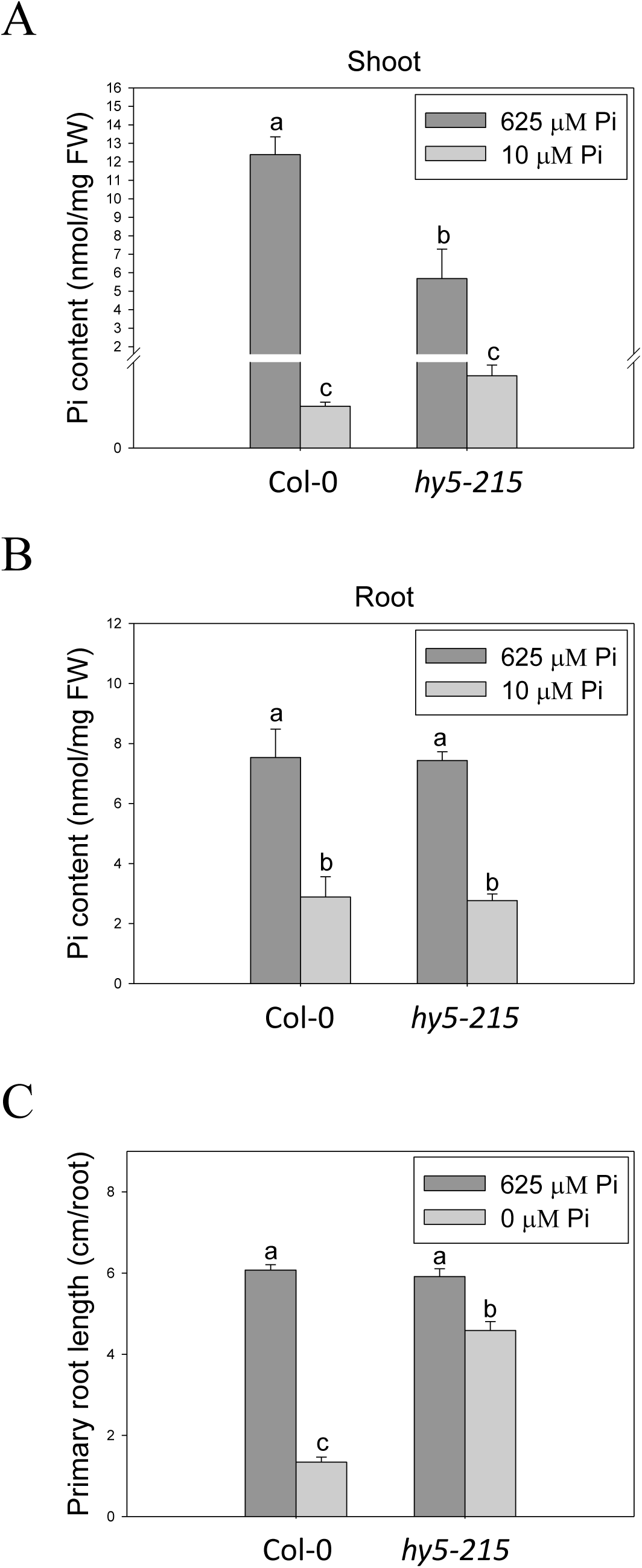
Pi content in wild-type and mutant seedlings in response to Pi treatment. (A)Soot Pi content in Col-0 and *hy5-215.* (B) Root Pi content in Col-0 and *hy5-215.* (C) PR length in Col-0 and *hy5-215* when Pi was sufficient or absent. The seedlings were grown on 1/2 MS medium with 625, 10 or 0 μM Pi for 10 days. Data represent means ± SE of four independent experiments. Different letters above the bars indicate statistically significant differences among the means based on ANOVA followed by Fisher’s LSD tests (*P* <0.05).

### Lower level of Pi deficiency-inducible membrane glycolipids in *hy5-215*

Since Pi deficiency tolerance of *hy5-215* was not due to Pi acquisition, we investigated Pi use efficiency in the mutant and wild type. Improvement of Pi utilization efficiency helps plants to conserve internal Pi and can involve the recycling of Pi from senescent tissues and the replacement of Pi from cellular structures or metabolic processes by alternative non-Pi compounds (Kobayashi et al. 2006, Rose et al. 2013). Membrane lipid remodeling, in which phospholipids are hydrolyzed and replaced by non-phosphorus glycolipids, such as sulfoquinovosyldiacylglycerol (SQDG) and digalactosyldiacylglycerol (DGDG), is a representative mechanism of Pi recycling, which improves Pi use efficiency (Kobayashi et al. 2006, Nakamura et al. 2013). Therefore, we analyzed the expression of genes involved in hydrolysis of phospholipids, novel phospholipase C gene (*NPC4*), and synthesis of SQDG and DGDG including *SQD1, SQD2, MGD2* and *MGD3* (monogalactosyldiacylglycerol synthetic genes) in the WT and *hy5-215.* All the analyzed genes were induced by Pi deficiency, but the expression levels were lower in *hy5-215* than in the WT (Supplementary Fig. S6A-E). The lipid composition calculated as the ratio of DGDG and PC (phosphatidylcholine), one of the major membrane phospholipids, is used as a marker to indicate a Pi-deficient state (Kobayashi et al. 2006). Enhancement of the DGDG/PC ratio represents an increase in DGDG biosynthesis to replace membrane phospholipids in response to Pi deficiency. A lower ratio of DGDG/PC was found in *hy5-215* under Pi-deficient conditions (Supplementary Fig. S6F), indicating that the increased tolerance to Pi deficiency in *hy5-215* mutants is not caused by increased free Pi from phospholipids.

### Identification of possible candidate genes responsible for Pi-deficiency tolerance in *hy5-215*

To determine the Pi-deficiency tolerance mechanism of *hy5-215*, we performed a transcriptome study using microarray. Consistent with previous reports, the well-known PSI genes were up-regulated in the WT under Pi deficiency. However, the expression levels of most PSI genes were significantly lower in *hy5-215*, including genes encoding high-affinity Pi transporters, ribonucleases, acid phosphatases, lipid remodeling and anthocyanin synthesis enzymes (Supplementary Table S1). Previously reported Pi deficiency-responsive TF genes in Arabidopsis mainly belong to the MYB and WRKY families (Rubio et al. 2001, Bustos et al. 2010, Yeh and Ohme-Takagi 2015). In this study, various TF genes, including *MYB, WRKY, AP2/ERF, bHLH, C2H2ZnF* and *MADS-box*, were up-regulated or down-regulated in *hy5-215* under Pi-deficient conditions (Supplementary Table S2), suggesting possible roles in the tolerance of *hy5-215* to Pi deficiency.

Liu et al. (wi2017) recently reported that HY5 negatively regulates expression of *PHR1* and its downstream PSI genes, and *hy5* mutant increases Pi and anthocyanin contents. According to their results, the longer root phenotype of *hy5* to phosphate starvation may result from the increased PSRs and Pi content. Although the root phenotypes of *hy5* are similar to our results, the expression of *PHR1* and PSI genes, and Pi and anthocyanin content were lower in the *hy5-215* mutant in our study (Fig. 3, Supplementary Fig. S2, S3, Table S1), which are consistent with previous reports that the expression of anthocyanin biosynthesis genes and anthocyanin accumulation are reduced in *hy5* (Lee et al. 2007, Jeong et al. 2010, Shin et al. 2013). Our results clearly show that the *hy5* tolerant phenotype to phosphate starvation is unlikely to be related to external Pi uptake because of similar growths of *hy5* on Pi-deficient and Pi-free conditions (Fig. 3C). Further information is required to address whether these inconsistencies result from different growth conditions and different plant tissues.

Unexpectedly, a significant number of photosynthesis-related and chlorophyll synthesis genes were down-regulated in roots but not shoots of *hy5-215* (Supplementary Fig. S7 and Table S3). Plant roots can accumulate chlorophyll and turn green under light illumination. The green roots are supposed to have photosynthetic ability as green leaves (Kobayashi et al. 2012). We therefore analyzed whether the Pi-deficiency tolerance of *hy5-215* is related to down-regulation of photosynthesis-related and chlorophyll synthesis genes, which may induce lower Pi consumption by decreasing photosynthesis in *hy5-215* roots. *GLK1* and *GLK2* have been shown to regulate expression of various photosynthetic genes in Arabidopsis roots (Kobayashi et al. 2012, Kobayashi et al. 2013). In addition, it was reported the roots of *35S:GLK1* accumulates much chlorophyll and are hypersensitive to Pi deficiency (Kang et al. 2012). We thus examined whether the *glk* mutants also show tolerance to Pi deficiency. The similar PR lengths between WT and *glk* mutants indicate *GLK1* and *GLK2* may not be involved in Pi-deficiency tolerance (Supplementary Fig. S8A, C). We further investigated the overexpression lines of *GLK1* and *GLK2* in *hy5-215* background *(35S:GLK1 hy5-215* and *35S:GLK2 hy5-215),* which have a recovered chlorophyll content as WT (Kobayashi et al. 2012). The *35S:GLK1 hy5-215* and *35S:GLK2 hy5-215* plants exhibited longer PR lengths under Pi deficiency similar to *hy5-215* (Supplementary Fig. S8B), suggesting that tolerance of *hy5-215* to Pi deficiency may not be related to chlorophyll content and photosynthetic activity.

To confirm this finding, the photosynthetic ability of *hy5-215* and WT plants was measured and compared, although photosynthetic gene expression in shoots was not significantly different between *hy5-215* and WT under Pi-sufficient or Pi-deficient conditions. As shown in Supplementary Fig. S9, the maximum quantum yield of photosystem II (Fv/Fm) and the actual quantum yield of photosystem II under light (YII) were reduced in the cotyledons of both WT and *hy5-215* in response to Pi deficiency. Although the measurement of Fv/Fm and Y_II_ of *hy5-215* under Pi sufficient treatment were lower than those of WT, there was no significant difference between WT and *hy5-215* in response to Pi depletion. In addition, Fv/Fm and Y_II_ in the true leaves of WT and *hy5-215* were not affected by our low Pi treatment. These data indicate that the tolerance of *hy5-215* to Pi deficiency is not related to photosynthetic ability (Supplementary Fig. S9). All together, these results indicate a novel mechanism other than the well-known PSRs may account for *hy5-215* tolerance to Pi deficiency.

### Light quality is involved in regulation of Pi deficiency response

Because HY5 acts as an integrator of different light signaling pathways downstream of multiple photoreceptor families and regulates photomorphogenesis (Cluis et al. 2004), we examined the effect of light on *hy5-215* tolerance to Pi deficiency. When the seedlings were grown in Pi-deficient conditions under continuous white light, WT and *hy5-215* PR lengths were 28% and 46% of PR lengths under Pi-sufficient conditions, respectively (Fig. 4A). Under continuous dark, there were no significant differences in PR growth between WT and *hy5-215* (Fig. 4B). These results, together with the results from long-day treatments (16 h light/8 h dark; Fig. 1B), indicate that increased light irradiation time inhibits Arabidopsis PR growth in Pi-deficient conditions. Therefore, light may play a role in *hy5-215* tolerance to Pi deficiency.

**Figure 4.**
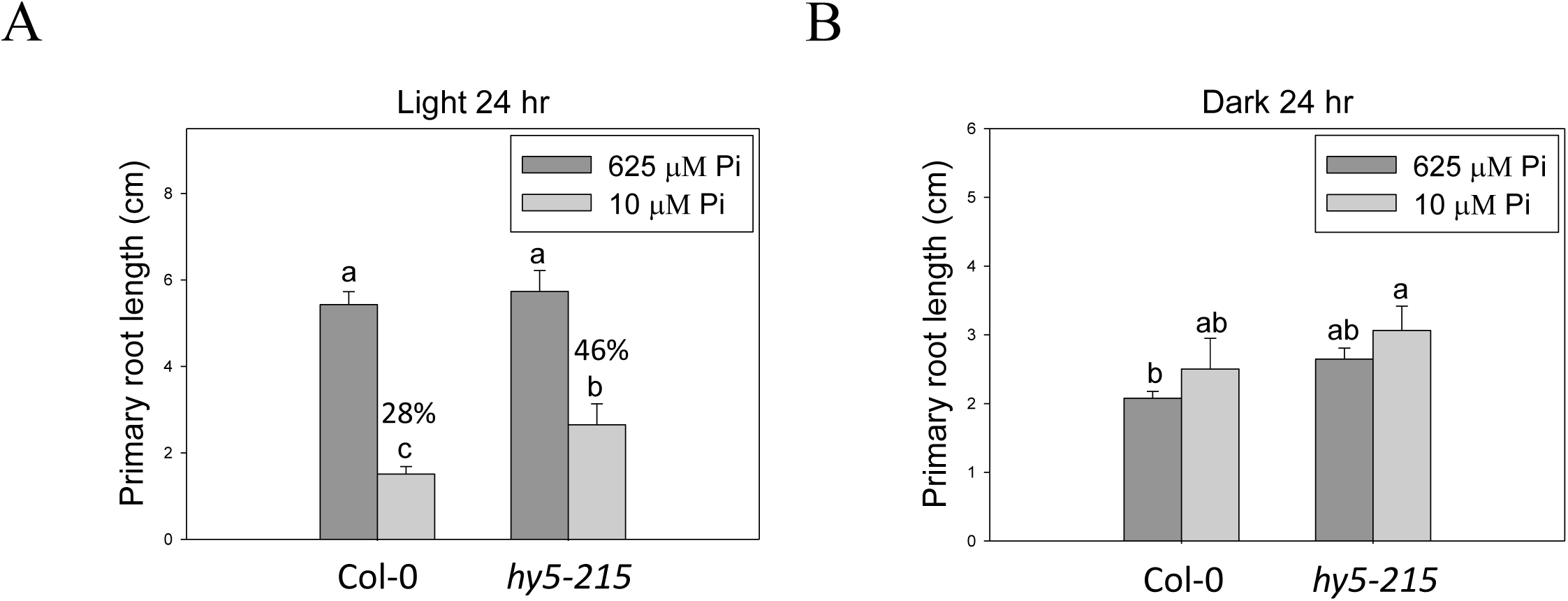
Effect of light on Pi-deficiency tolerance in Arabidopsis. The seedlings were grown on 1/2 MS media with 625 or 10 μM Pi under continuous light (A) or dark (B) treatments. The PR length was measured after 10 days of growth. Data represent means ± SE of four independent experiments. Different letters above the bars indicate statistically significant differences among the means based on ANOVA followed by Fisher’s LSD test (*P* <0.05).

To better understand light effects on Pi-deficiency tolerance, Arabidopsis plants were grown under continuous blue (B), red (R) and far-red (FR) light. PR growth was inhibited by Pi deficiency in the WT under continuous B light to a similar extent as was observed in white light. In contrast, the same level of inhibition by Pi deficiency under B light was not observed in *hy5-215* (Fig. 5A). Interestingly, PR growth was not inhibited by Pi deficiency in both WT and *hy5-215* when grown under continuous R and FR irradiation (Fig. 5B-D and Supplementary Fig. S10). These results indicate that the tolerance of *hy5-215* to Pi deficiency is negatively regulated by B light and is not related to R and FR light. To further confirm this finding, the B light receptor mutants, *cry1 cry2* and *phot1 phot2*, were examined under Pi deficiency. Indeed, a tolerant phenotype to Pi deficiency was found in these two mutants (Fig. 5E-F). Therefore, the tolerance of *hy5-215* to Pi deficiency likely results from blockage of B light responses, and the tolerance mechanism may be related to enhancement of internal Pi recycling or utilization efficiency but not external Pi acquisition due to the tolerant phenotype of *hy5-215* under Pi-free condition. Our findings may provide valuable insights for developing Pi deficiency-tolerant crops in the future. Furthermore, light quality-regulated responses to Pi deficiency may allow indoor plant growers to reduce Pi fertilizer application through proper illumination.

**Figure 5.**
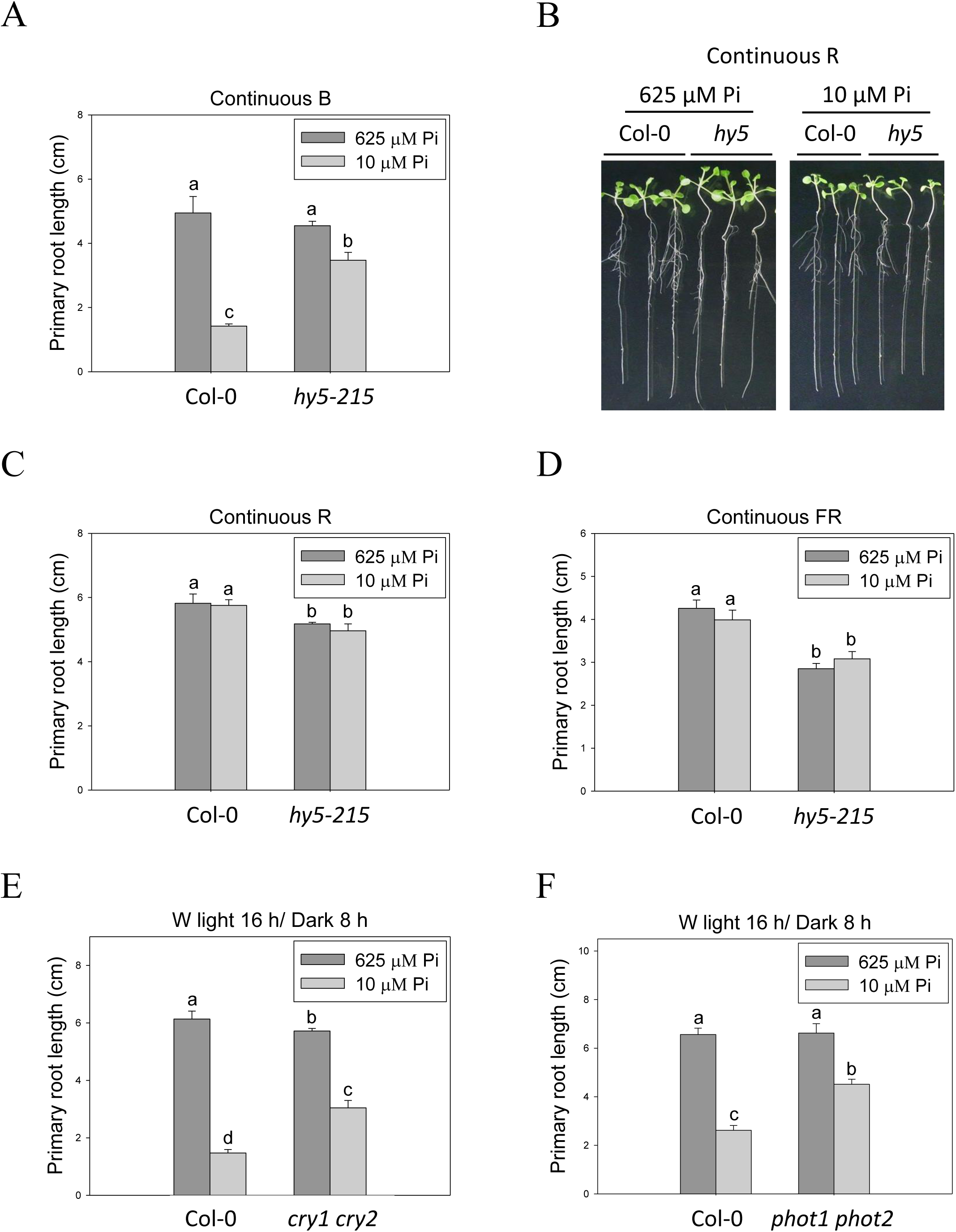
Effect of light quality on primary root length in Arabidopsis. The seedlings were grown on 1/2 MS media with 625 or 10 μM Pi under continuous blue (B), red (R) or far red (FR) light treatments, respectively (A-D). The blue light receptor mutants, *cry1 cry2* (E) and *phot1 phot2* (F), were grown on Pi-sufficient and Pi-deficient media under long-day condition (16 h light/8 h dark). PR length was measured after 10 days of growth. Data represent means ± SE of four independent experiments. Different letters above the bars indicate statistically significant differences among the means based on ANOVA followed by Fisher’s LSD test (*P* <0.05).

## Materials and Methods

### Plant materials and growth conditions

The surface-sterilized seeds of *Arabidopsis thaliana* wild type [ecotypes Columbia (Col-0)] and mutants (*hy5-215, slr-1, hy5-215 slr-1, glk1, glk2, glk1 glk2, cry1 cry2, phot1 phot2*), and transformants *(35S:GLK1 hy5-215* and *35S:GLK2 hy5-215)* were sown on 1/2 Murashige and Skoog (MS) agar plates containing 625 μM KH_2_PO_4_ (Pi sufficient) or 10 μM KH_2_PO_4_ (Pi deficient). Each experiment used 10 plants and was replicated three to four times. The seedlings were grown at 22°C and illuminated with 100-125 μmol m^-2^ s^-1^ white light for 16 hours per day or with blue (B), red (R) and far-red (FR) light for 24 hours. For determination of primary root (PR) length and fresh weight, the seedlings were cultured on vertical and horizontal plates for 10 and 14 days, respectively. The seedlings were then collected for photographs, measurement of PR length and fresh weight, and further experiments.

### Quantification of anthocyanin content

The shoots of 10-day-old seedlings were frozen in liquid nitrogen, ground into a powder, and then re-suspended in an extraction buffer containing 45% methanol and 5% acetic acid. The supernatant was taken after centrifugation at 12,000 rpm for 10 minutes. Anthocyanin content was calculated by the absorbance at 530 and 637 nm as described previously (Matsui et al. 2004).

### Determination of acid phosphatase activity

The histochemical staining of acid phosphatase activity was performed according to the method described by Yu et al. (2012) with some modifications. The roots of 10-day-old seedlings were overlaid with a 0.1% agar solution containing 0.01% 5-bromo-4-chloro-3-indolyl phosphate (BCIP). The acid phosphatase activity indicated by blue color on the root surface was observed and photographed after 6 to 24 hours.

### Determination of lipid composition

Seedlings were grown on 1/2 MS medium with 625 μM Pi for 10 days and then transferred to 1/2 MS medium with 625 μM Pi or 10 μM Pi for 10 days. Samples were collected and immediately frozen in liquid nitrogen. Lipids were then extracted and analyzed by the method described by Kobayashi et al. (2006).

### RNA isolation, reverse-transcription quantitative PCR (RT-qPCR), and microarray analyses

Total RNA was extracted by using the RNeasy Plant Mini kit (QIAGEN, Hilden, Germany) following the manufacturer’s instructions. One μg of total RNA was subjected to first-strand cDNA synthesis using the PrimeScript RT reagent kit (Takara). Quantitative RT-qPCR was performed by the SYBR green method using the ABI7300 real-time PCR system (Applied Biosystems) as described previously (Mitsuda et al. 2005). The *UBQ1* gene was used as an internal control. The microarray experiments and the data analysis were conducted by the method described by Mitsuda et al. (2005). Three or four biological replicates were included in each experiment.

### Measurement of photosynthetic activity

The maximum quantum yield of photosystem II (F_v_/F_m_) and actual quantum yield of photosystem II in light (YII) of cotyledons and true leaves were measured according to the method described by Kobayashi et al. (2013).

### Statistical analysis

All the experiments were performed in a completely randomized design. Data on root length (cm) and seedling fresh weight (mg) were recorded after growth for 10 and 14 days, respectively. Analysis of variance (ANOVA) and mean comparisons using least significant difference (LSD) tests were conducted. Data represent means of three or four independent experiments. Different letters above bars indicate statistically significant differences (*P* <0.05).

### Accession numbers

Arabidopsis Genome Initiative numbers described in this article are as follows: *ACP5* (At3g17790), *CHS* (At5g13930), *DFR* (At5g42800), *GLK1* (At2g20570), *GLK2* (At5g44190), *HY5* (At5g11260), *LDOX* (At4g22880), *MGD2* (At5g20410), *MGD3* (At2g11810), *MYB75* (At1g56650), *MYB90* (At1g66390), *NPC4* (At3g03530), *PHT1;2* (At5g43370), *PHT1;3* (At5g43360), *PHT1;4* (At2g38940), *PHT1;5* (At2g32830), *PHT1;7* (At3g54700), *PHT1;8* (At1g20860), *PHT1;9* (At1g76430), *RNS1* (At2g02990), *SLR/IAA14* (At4g14550), *SQD1* (At4g33030), *SQD2* (At5g01220) and *UF3GT* (AT5G54060).

## Abbreviations

B: blue
DGDG: digalactosyldiacylglycerol
FR: far-red
hps: hypersensitive to phosphate starvation
HY5: ELONGATED HYPOCOTYL 5
LR: lateral root
MGD: monogalactosyldiacylglycerol
NPC4: novel phospholipase C
PC: phosphatidylcholine
PHL1: PHR1-like 1
PHO1: PHOSPHATE1
PHR1: PHOSPHATE STARVATION RESPONSE 1
Pi: inorganic phosphate
PR: primary root
PSI: phosphate starvation-induced
PSR: phosphate starvation response
R: red
slr-1: solitary-root-1
SQDG: sulfoquinovosyldiacylglycerol
TF: transcription factor
WT: wild type

## Acknowledgements

We thank Professor Kiyotaka Okada (National Institute for Basic Biology, Japan) for providing *hy5-215* seeds, Professor Shigo Takagi (Osaka University, Japan) and Professor Hirokazu Tsukaya (The University of Tokyo, Japan) for *phot1, phot2* and *phot1 phot2* seeds, Professor Christian Fankhauser (University of Lausanne, Switzerland) for *cry1 cry2* seeds and Dr. Yukari Nagatoshi (JIRCAS, Japan) for *glk1, glk2* and *glk1 glk2* seeds. We also thank Dr. Sumire Fujiwara and Dr. Yukari Nagatoshi for discussion on the research, and Naomi Ujiie, Machiko Onuki, Yukie Kimura and Sumiko Takahash for technical assistance. This work was supported by grants from The Japan Society for the Promotion of Science (JSPS) and The Japan Science and Technology Agency (JST).

## References

Bari R., Datt Pant B., Stitt M., Scheible W.R. (2006) PHO2, microRNA399, and PHR1 define a phosphate-signaling pathway in plants. Plant Physiol. 141: 988-999.

Bayle V., Arrighi J.F., Creff A., Nespoulous C., Vialaret J., Rossignol M. et al. (2011) Arabidopsis thaliana high-affinity phosphate transporters exhibit multiple levels of posttranslational regulation. Plant Cell 23: 1523-1535.

Bustos R., Castrillo G., Linhares F., Puga M.I., Rubio V., Pérez-Pérez J. et al. (2010) A central regulatory system largely controls transcriptional activation and repression responses to phosphate starvation in Arabidopsis. PLoS Genet. 6: e1001102.

Chen Z.H., Nimmo G.A., Jenkins G.I., Nimmo H.G. (2007) BHLH32 modulates several biochemical and morphological processes that respond to Pi starvation in *Arabidopsis*. Biochem. J. 405: 191-198.

Chen Y.F., Li L.Q., Xu Q., Kong Y.H., Wang H., Wu W.H. (2009) The WRKY6 transcription factor modulates *PHOSPHATE1* expression in response to low Pi stress in *Arabidopsis*. Plant Cell 21: 3554-3566.

Chiou T.J., Aung K., Lin S.I., Wu C.C., Chiang S.F., Su C.L. (2006) Regulation of phosphate homeostasis by microRNA in Arabidopsis. Plant Cell 18: 412-421.

Cluis C.P., Mouchel, C.F., Hardtke C.S. (2004) The Arabidopsis transcription factor HY5 integrates light and hormone signaling pathways. Plant J. 38: 332-347.

Devaiah B.N., Karthikeyan A.S., Raghothama K.G. (2007a) WRKY75 transcription factor is a modulator of phosphate acquisition and root development in Arabidopsis. Plant Physiol. 143: 1789-1801.

Devaiah B.N., Nagarajan V.K., Raghothama K.G. (2007b) Phosphate homeostasis and root development in Arabidopsis are synchronized by the zinc finger transcription factor ZAT6. Plant Physiol. 145: 147-159.

Devaiah B.N., Madhuvanthi R., Karthikeyan A.S., Raghothama K.G. (2009) Phosphate starvation responses and gibberellic acid biosynthesis are regulated by the *MYB62* transcription factor in *Arabidopsis*. Mol. Plant 2: 43-58.

Fukaki H., Tameda S., Masuda H., Tasaka M. (2002) Lateral root formation is blocked by a gain-of-function mutation in the *SOLITARY-ROOT/IAA14* gene of Arabidopsis. Plant J. 29: 153-168.

Gaxiola R.A., Edwards M., Elser J.J. (2011) A transgenic approach to enhance phosphorus use efficiency in crops as part of a comprehensive strategy for sustainable agriculture. Chemosphere 84: 840-845.

Gilbert N. (2009) Environment: The disappearing nutrient. Nature 461: 716-718.

Gruber B.D., Giehl R.F., Friedel S., von Wirén N. (2013) Plasticity of the Arabidopsis root system under nutrient deficiencies. Plant Physiol. 163: 161-179.

Hamburger D., Rezzonico E., MacDonald-Comber Petétot J., Somerville C., Poirier Y. (2002) Identification and characterization of the *Arabidopsis PHO1* gene involved in phosphate loading to the xylem. Plant Cell. 14: 889-902.

Jeong S.W., Das P.K., Jeoung S.C., Song J.Y., Lee H.K., Kim Y.K. et al. (2010) Ethylene suppression of sugar-induced anthocyanin pigmentation in Arabidopsis. Plant Physiol. 154: 1514-1531.

Kang J., Yu H., Tian C., Zhou W., Li C., Jiao Y. et al. (2014) Suppression of photosynthetic gene expression in roots is required for sustained root growth under phosphate deficiency. Plant Physiol. 165: 1156-1170.

Kobayashi K., Masuda T., Takamiya K., Ohta H. (2006) Membrane lipid alteration during phosphate starvation is regulated by phosphate signaling and auxin/cytokinin cross-talk. Plant J. 47: 238-248.

Kobayashi K., Baba S., Obayashi T., Sato M., Toyooka K., Keränen M. et al. (2012) Regulation of root greening by light and auxin/cytokinin signaling in Arabidopsis. Plant: Cell 24: 1081-1095.

Kobayashi K., Sasaki D., Noguchi K., Fujinuma D., Komatsu H., Kobayashi M. et al. (2013) Photosynthesis of root chloroplasts developed in Arabidopsis lines overexpressing *GOLDEN2-LIKE* transcription factors. Plant Cell Physiol. 54: 1365-1377.

Lee J., He K., Stolc V., Lee H., Figueroa P., Gao Y. et al. (2007) Analysis of transcription factor HY5 genomic binding sites revealed its hierarchical role in light regulation of development. Plant Cell 19: 731-749.

Lei M., Liu Y., Zhang B., Zhao Y., Wang X., Zhou Y. (2011) Genetic and genomic evidence that sucrose is a global regulator of plant responses to phosphate starvation in Arabidopsis. Plant Physiol. 156: 1116-1130.

Liu F., Wang Z., Ren H., Shen C., Li Y., Ling H.Q. et al. (2010) OsSPX1 suppresses the function of OsPHR2 in the regulation of expression of *OsPT2* and phosphate homeostasis in shoots of rice. Plant J. 62: 508-517.

Liu Y., Xie Y., Wang H., Ma X., Yao W., Wang H. (2017) Light and Ethylene Coordinately Regulate the Phosphate Starvation Response through Transcriptional Regulation of *PHOSPHATE STARVATION RESPONSE1*. Plant Cell 29: 2269-2284.

Matsui K., Tanaka H., Ohme-Takagi M. (2004) Suppression of the biosynthesis of proanthocyanidin in Arabidopsis by a chimeric PAP1 repressor. Plant Biotechnol. J. 2: 487-493.

Misson J., Raghothama K.G., Jain A., Jouhet J., Block M.A., Bligny R. et al. (2005) A genome-wide transcriptional analysis using Arabidopsis thaliana Affymetrix gene chips determined plant responses to phosphate deprivation. Proc. Natl. Acad. Sci. USA. 102: 11934-11939.

Mitsuda N., Seki M., Shinozaki K., Ohme-Takagi M. (2005) The NAC transcription factors NST1 and NST2 of Arabidopsis regulate secondary wall thickenings and are required for anther dehiscence. Plant Cell 17: 2993-3006.

Nagarajan V.K., Jain A., Poling M.D., Lewis A.J., Raghothama K.G., Smith A.P. (2011) Arabidopsis Pht 1 ;5 mobilizes phosphate between source and sink organs, and influences the interaction between phosphate homeostasis and ethylene signaling. Plant Physiol. 156: 1149-1163.

Nakamura Y. (2013) Phosphate starvation and membrane lipid remodeling in seed plants. Prog. Lipid Res. 52: 43-50.

Niu Y.F. (2013) Responses of root architecture development to low phosphorus availability: a review. Ann. Bot. 112: 391-408.

Nussaume L., Kanno S., Javot H., Marin E., Pochon N., Ayadi A., et al. (2011) Phosphate Import in Plants: Focus on the PHT1 Transporters. Front. Plant Sci. 2: 83.

Oyama T., Shimura Y., Okada K. (1997) The *Arabidopsis HY5* gene encodes a bZIP protein that regulates stimulus-induced development of root and hypocotyl. Genes Dev. 11: 2983-2995.

Péret B., Clément M., Nussaume L., Desnos T. (2011) Root developmental adaptation to phosphate starvation: better safe than sorry. Trends Plant Sci. 16: 442-450.

Pérez-Torres C.A., López-Bucio J., Cruz-Ramírez A., Ibarra-Laclette E., Dharmasiri S., Estelle M. et al. (2008) Phosphate availability alters lateral root development in Arabidopsis by modulating auxin sensitivity via a mechanism involving the TIR1 auxin receptor. Plant Cell 20: 3258-3272.

Plaxton W.C., Tran H.T. (2011) Metabolic adaptations of phosphate-starved plants. Plant Physiol. 156: 1006-1015.

Poirier Y., Bucher M. (2002) Phosphate transport and homeostasis in Arabidopsis. In: Somerville CR, Meyerowitz, EM, editors. The Arabidopsis Book e0024.

Raghothama K.G. (2000) Phosphate transport and signaling. Curr. Opin. Plant Biol. 3: 182-187.

Rose T.J., Liu L., Wissuwa M. (2013) Improving phosphorus efficiency in cereal crops: Is breeding for reduced grain phosphorus concentration part of the solution? Front. Plant Sci. 4: 444.

Rubio V., Linhares F., Solano R., Martín A.C., Iglesias J., Leyva A. et al. (2001) A conserved MYB transcription factor involved in phosphate starvation signaling both in vascular plants and in unicellular algae. Genes Dev. 15: 2122-2133.

Sato A., Miura K. (2011) Root architecture remodeling induced by phosphate starvation. Plant Signal. Behav. 6: 1122-1126.

Shin D.H., Choi M., Kim K., Bang G., Cho M., Choi S.B. et al. (2013) HY5 regulates anthocyanin biosynthesis by inducing the transcriptional activation of the MYB75/PAP1 transcription factor in Arabidopsis. FEBS Lett. 587: 1543-1547.

Thibaud M.C., Arrighi J.F., Bayle V., Chiarenza S., Creff A., Bustos R., et al. (2010) Dissection of local and systemic transcriptional responses to phosphate starvation in Arabidopsis. Plant J. 64: 775-789.

Ticconi C.A., Abel S. (2004) Short on phosphate: plant surveillance and countermeasures. Trends Plant Sci. 9: 548-555.

Wissuwa M. (2003) How do plants achieve tolerance to phosphorus deficiency? Small causes with big effects. Plant Physiol. 133: 1947-1958.

Woo J., MacPherson C.R., Liu J., Wang H., Kiba T., Hannah M.A. et al. (2012) The response and recovery of the *Arabidopsis thaliana* transcriptome to phosphate starvation. BMC Plant Biol. 12: 62.

Wu P., Ma L., Hou X., Wang M., Wu Y., Liu F. et al. (2003) Phosphate starvation triggers distinct alterations of genome expression in Arabidopsis roots and leaves. Plant Physiol. 132:1260-1271.

Yeh C.M., Ohme-Takagi M. (2015) Transcription factors involved in acid stress responses in plants. Nucleus 58: 191-197.

Yeh C.M., Ohme-Takagi M., Tsai W.C. (2017) Current understanding on the roles of ethylene in plant responses to phosphate deficiency. Int. J. Plant Biol. Res. 5: 1058.

Yu H., Luo N., Sun L., Liu D. (2012) HPS4/SABRE regulates plant responses to phosphate starvation through antagonistic interaction with ethylene signalling. J. Exp. Bot. 63: 4527-4538.

Zhou J., Jiao F., Wu Z., Li Y., Wang X., He X. et al. (2008) *OsPHR2* is involved in phosphate-starvation signaling and excessive phosphate accumulation in shoots of plants. Plant Physiol. 146: 1673-1686.

